# Lactate receptor, HCAR1, in neonatal hypoxic-ischemic encephalopathy

**DOI:** 10.1101/2025.09.16.676275

**Authors:** Jennifer Burnsed, Angelina June, Maria Marlicz, Emmy Zucker, Ellie Kain-Kuzienski, Hannah Mulhern, Leanne Maharaj, Chengsan Sun, Suchitra Joshi, Huayu Sun, Jaideep Kapur

**Author notes:** Corresponding author: Jennifer Burnsed, MD, MS, Associate Professor of Pediatrics, Division of Neonatology, University of Virginia.

## Abstract

**Introduction:** Hydroxycarboxylic acid receptor 1 (HCAR1) is a G-protein coupled receptor for lactate that is expressed in the brain and plays a role in neuronal excitability, angiogenesis, and repair after injury. Hypoxic-ischemic encephalopathy (HIE) is the most common cause of brain injury and seizures in term neonates. The goal of this study was to describe HCAR1 expression and function in the neonatal brain and further understand its role in HIE.

**Methods:** HCAR1 expression was measured using qRT-PCR in developing mice (postnatal day (p)10, 20, 30, 50). Electrophysiology was used to measure neuronal properties and spontaneous excitatory postsynaptic currents (sEPSC) in hippocampal principal neurons from HCAR1 knockout and wildtype mice when exposed to lactate. Then, p10 HCAR1 knockout and wildtype mice were exposed to hypoxia-ischemia (HI) and placed on electroencephalography (EEG) to compare seizure burden. HCAR1 expression after neonatal HI was measured with PCR.

**Results:** HCAR1 is expressed at p10 at similar levels to adults (n=6/group; 1-way ANOVA, p<0.0004). Lactate decreases amplitudes and sEPSC frequency in wildtype (p<0.0001, p<0.001) but not HCAR1 knockout mice (p<0.26, p=0.91). After HI, HCAR1 knockout mice have higher seizure burden (2318s in WT vs. 6497s in KO (p=0.04)) and behavioral seizure scores (4 in WT vs. 6 in KO (p<0.001)) than wildtypes. HCAR1 expression increased 24h post-HI but drops to below baseline at 48h post-HI. (n=3/group; p=0.02).

**Conclusion:** HCAR1 is expressed on neurons in the developing mouse brain. Lactate decreases neuronal excitability via HCAR1. HCAR1 is upregulated post-HI and mice lacking HCAR1 exhibit worse neonatal HI seizures.

**Highlights:**

- HCAR1, a G-protein coupled, lactate receptor is expressed in the developing mouse brain
- HCAR1 is expressed on neurons in the neonatal mouse hippocampus
- HCAR1 is upregulated 24 hours after neonatal hypoxia-ischemia
- Mice lacking HCAR1, exhibit worse neonatal hypoxic-ischemic seizures

## Introduction

Hydroxycarboxylic acid receptor 1 (HCAR1) is a G-protein coupled receptor for lactate that is expressed in the brain, adipose tissue, skeletal muscle, kidney, liver, pancreas, and various immune cells^1–3^. In the brain, HCAR1 has been described to play a role in neuronal excitability, angiogenesis, neural progenitor cell development, and repair after injury^4–10^.

Hypoxic-ischemic encephalopathy is the most common cause of brain injury and seizures in full- and near-term infants^11,12^. Currently, the only neuroprotective treatment in this population is therapeutic hypothermia (TH), which only improves outcomes in about a third of patients^13–15^. New and adjuvant treatments are needed in this population to improve outcomes.

In preclinical studies, lactate administered to neonatal rodents after hypoxia-ischemia (HI) reduced lesion volume and improved performance on memory and sensorimotor testing^16,17^. Further, rodents lacking HCAR1 exhibit increased infarct size and reduced tissue regeneration after HI^7,18^.

Understanding of how HCAR1 differs in the developing brain and its specific role in injury and neuroprotection is lacking. The goal of this study was to describe HCAR1 expression and function in the neonatal brain and further understand its role in HI injury.

## Methods and materials

### Animals

All animals used in this study were handled according to a University of Virginia Animal Care and Use Committee approved protocol (#4071). Housing was in accordance with the National Institutes of Health Guide for Care and Use of Laboratory Animals (US Department of Health and Human Services 85-23, 2011). C57Bl/6 mice were used for the qPCR and RNAscope portions of this study. We used HCAR1 global knockout mice for all remaining experiments in this study, previously described^10^.

### qPCR for HCAR1

To measure HCAR1, we used a qRT-PCR reaction. Brains from mice at several different ages (p10, 20, 30 and 50) were isolated to measure and compare normal developmental levels of HCAR1 mRNA. Brains were also obtained from mice exposed to HI at three timepoints: immediately following HI, 24 hours after HI and 48 hours after HI. The hemisphere ipsilateral to the carotid ligation was used for these experiments. Total RNA isolation and cDNA synthesis were performed^19^ and the expression of HCAR1 was measured using sensiFAST SYBR fluorescein kit (BioLine) using the primer targeting the HCAR1 gene. Specificity of primers had been previously tested and validated as described in prior studies^10^. Samples were all run in replicate and negative controls were run for each gene to confirm primer specificity. Relative HCAR1 mRNA expression was quantified using the comparative CT methods and expressed as fold change using the ΔΔCT formula^20^.

### RNAscope

In order to examine HCAR1 expression in the naive p10 mouse brain we used RNAscope for HCAR1 with NeuN co-staining to label neurons. P10 mice were deeply anesthetized using isoflurane and transcardially perfused using 4% paraformaldehyde.

Brains were fixed in 4% PFA at 4°C overnight and sliced at 15μm thickness on a vibratome. Slices were mounted on charged slides in preparation for RNAscope and immunohistochemistry using fixation and permeabilization steps that optimize preservation of both RNA and protein structures. All sections for each experimental run were mounted and reacted on the same slide to control for variation in experimental conditions and solutions. Slices were rinsed in sterile water and incubated with “pretreat4” from the RNAscope Multiplex Fluorescent Assay kit (Advanced Cell Diagnostics) for 30 minutes at 40°C. Sections were counterstained with DAPI. In accordance with manufacturer protocol for the RNAscope Multiplex Fluorescent Detection Kit v2 (Advanced Cell Diagnostics, Cat#323110), sections were incubated with RNA scope catalog oligonucleotide probes for Mus musculus G protein-coupled receptor 81 (GPR81, AKA HCAR1), negative control probe (DAPB, bacillus subtilis strain SMY methylglyoxal synthase, mgsA, partial cds dihydrodipicolinate reductase, dapB gene, complete cds and YpjD gene, partial cds; Cat# 317421), and positive control probe (Mm-Polr2a, Mus musculus polymerase (RNA) II (DNA directed) polypeptideA (Polr2a, mRNA; Cat#312471). For immunohistochemistry, after target retrieval steps, the sections were incubated in the primary antibody overnight at 4ºC (neuronal marker, anti-NeuN). Since we used three fluorophores (to visualize RNA, protein, and DAPI for nuclear stain), we scanned each fluorophore separately to prevent signal bleed through. We optimized the RNAscope technique to reliably visualize the HCAR1 transcript using positive and negative controls, and HCAR1 KO mice (a second negative control).

### Electrophysiology

#### Acute slice preparation

p10 HCAR1 knockout and wildtype littermates were deeply anesthetized using Halothane and then decapitated. Brains were quickly removed and immersed in oxygenated (95% O_2_/5%CO_2_) ice-cold (0-4°C) slicing buffer (in mM 65.5 NaCl, 2 KCl, 5 MgSO_4_, 25 NaHCO_3_, 1.1 KH_2_PO_4_, 1 CaCl_2_, 10 glucose, and 113 sucrose; 300 mOsm, pH 7.35-7.45). Slices cut from the hippocampus (300-350µm thick) were prepared using a Vibratome (Leica VT1200S, Germany) to obtain CA1. The slices were then transferred to an incubation chamber, where they were kept submerged at 33°C in artificial CSF (aCSF, containing (in mM), 127 NaCl, 2 KCl, 1.5 MgSO_4_, 25.7 NaHCO_3_, 1.5 CaCl_2_, and 10 glucose; 300 mOsm, bubbled with 95% O_2_/5%CO_2_, pH 7.35-7.45).

#### Patch-clamp recordings

CA1 neurons were identified under the differential interface contrast (DIC) microscope and recorded. During the recording, slices were transferred to a recording chamber perfused with aCSF at 2-3 mL/min. Recordings were performed by using a Multiclamp 700A amplifier (Molecular Devices, Union City, CA). The data were collected using pClamp 10.2 software (Molecular Devices), filtered at 2kHz, sampled at 10KHz (Digidata 1440A; Molecular Devices), and stored on a Pentium-based computer. Recording pipettes were fabricated from borosilicate glass (1.5mm outer diameter, 0.86mm inner diameter; Sutter Instruments, Novato, CA) using a Flaming-Brown micropipette puller (P-1000; Sutter Instruments, Novato, CA). EPSCs were recorded in voltage-clamp mode. Tight-seal whole-cell recordings were obtained using standard techniques^21^. Recording pipettes had open-tip resistances of 6-8MΩ) and were filled with the internal solution (containing (in mM): 110 D-gluconic acid, 110 CsOH, 10 CsCl, 1 EGTA, 1 CaCl2, 10 HEPES, 5 MgATP, 5 lidocaine, pH 7.3; 290 mOsm). Electrode capacitance was compensated electronically and access resistance was monitored continuously. The recording was terminated if the series resistance increased by 20%. CA1 cells were identified visually, patched in voltage-clamp configuration, and recorded in a holding potential of −65mV for EPSCs. Current-clamp recordings were performed to measure cell membrane properties. The recordings were performed in tight-seal (seal ≥1GigΩ) current-clamp mode. The internal solution consisted of (in mM): 135 K-gluconate, 7 KCl, 10 HEPES, 0.5 EGTA, 2.5 NaCl, 4 Mg-ATP and 0.3 NaGTP.

We recorded sEPSCs from CA1 principal neurons and after a 5-minute baseline recording, we applied 3mM of sodium lactate for 10 minutes.

### HI model

HI injury was modeled in postnatal day (p)10 mice with permanent unilateral left carotid artery ligation followed by 1 hour of recovery and then 1 hour of hypoxia at FiO2 = 0.8, as previously described^22,23^. Mice were anesthetized using isoflurane during carotid ligation. The surgical field and hypoxia/recording chamber were on warming pads and temperature was controlled to maintain normothermia.

### Electrode implantation and continuous video EEG monitoring

In order to characterize acute seizure activity and behavior, mice had continuous video EEG recording throughout the experimental period as previously described^24–26^. On p9, 24 hours prior to HI, unipolar insulated stainless steel depth electrodes (0.005 inches bare diameter, 0.008 inches coated; A-M Systems, Sequim, WA, USA) were stereotactically implanted in the bilateral parietal cortex (−1.22mm Dorsal-Ventral (DV), ± 0.5mm Medial-Lateral (ML), −1.75mm Deep (D)) along with reference electrodes in the cerebellum. Mice were anesthetized with isoflurane during electrode implantation. After 24 hours of recovery, mice were exposed to HI as described above. A unity gain impedance matching headstage (TLC2274 Quad Low-Noise Rail-to-Rail Operational Amplifier; Texas Instruments, Dallas, TX, USA) was used for recordings. Baseline EEG recording began 30 minutes prior to carotid ligation, mice were briefly removed from EEG for carotid ligation, then placed back in the recording chamber after ligation. Recording continued through ligation recovery for 1 hour, through the hour of hypoxia, and reoxygenation period of 30 minutes^24^. This equates to three hours of total video EEG recording per animal.

### Data Analysis

#### PCR

Relative HCAR1 mRNA expression was quantified using the comparative CT methods and expressed as fold change using the ΔΔCT formula^20^.

#### Electrophysiology

Amplitude and frequency of sEPSCs were analyzed by MiniAnalysis software. Decay kinetics were evaluated using weighted tau^27^. Current clamp data were analyzed using Clampex 10.5 and Minianalysis. Student paired t-test was used for cells recorded in the same slice (before and after lactate application). As noted in the results, the unpaired t-test, one-way ANOVA, and two-way ANOVA were also used when appropriate. For the sEPSC analysis, we used Kolmogorov-Smirnov since the results were not normally distributed. p-values <0.05 were considered significant.

#### EEG

LabChart Pro (ADInstruments, Colorado Springs, CO, USA) was used to collect and analyze video EEG data. Video EEG was reviewed and marked for seizures by a researcher (A.J.) blinded to the group and validated with randomly excerpted segments marked by a second blinded researcher (J.B.). Reviews were compared to confirm agreement. A behavioral seizure score (BSS) adapted for neonatal rodents was used, as previously described^28^. GraphPad Prism 9.5.0 (Dotmatrics, San Diego, CA, USA) was used for statistical analysis. Descriptive statistics were reported as either mean ± standard deviation or median ± standard deviation. Mean seizure time, total seizure burden (number of seizures x time per seizure) and BSS median was reported. Criteria for electrographic seizure was as previously described^29,30^.

## Results

### HCAR1 is similarly expressed in neonatal and adult mice

HCAR1 mRNA was measured using qPCR at multiple developmental timepoints including p10, p20, p30, p50 (Figure 1A; n=6/group; 1-way ANOVA, p<0.0004). Post-hoc multiple comparisons test (Tukey’s multiple comparisons test) showed that HCAR1 expression in p10 mice is similar to p30 (p=0.96) and p50 (p=0.19). However, mice from the p20 group had significantly lower HCAR1 expression compared to p10 (p=0.019) and p50 (p=0.0002).

**Figure 1:**
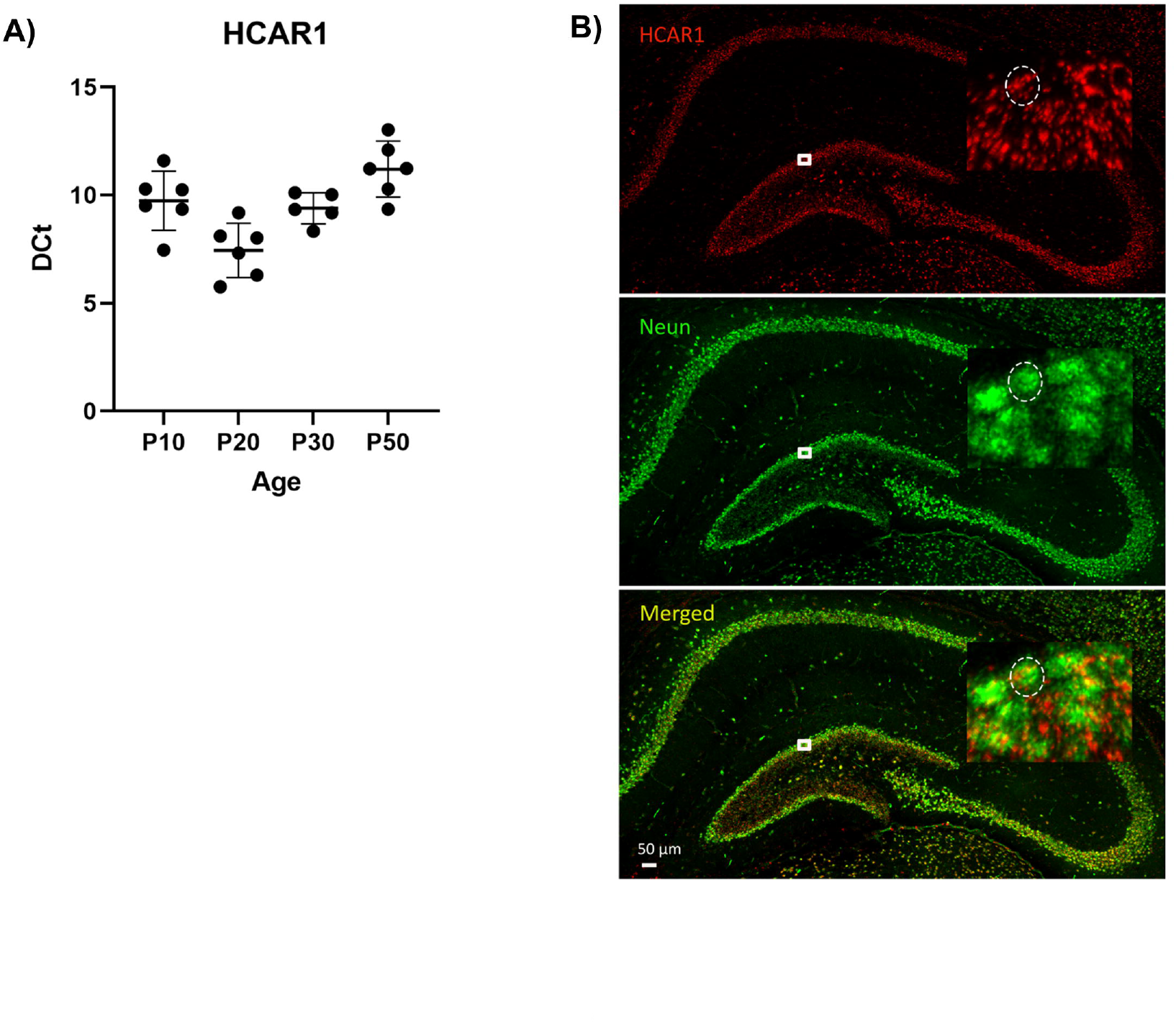
HCAR1 is expressed in the neonatal mouse brain at levels similar to adults. **A)** qPCR at multiple developmental timepoints shows that HCAR1 mRNA expression in the p10 mouse brain is similar to p30 and p50. (n=6/group; 1-way ANOVA, p<0.0004; multiple comparisons showed no significant difference between p10 and p30 or p50, p=0.96 and 0.19 respectively); **B)** HCAR1 expression in hippocampal neurons of the p10 mouse brain. Representative confocal images of coronal hippocampal sections of HCAR1 RNAscope (red) and NeuN (a neuronal marker) immunostaining (green). (Top) HCAR1 expression is observed predominantly in the granule cell layer and pyramidal cell layer of the hippocampus and displaying a punctate distribution pattern. (Middle) NeuN immunoreactivity highlights neuronal nuclei within the same region. (Bottom) Merged image demonstrates colocalization (yellow) of HCAR1 with NeuN, suggesting neuronal expression of HCAR1. Insets in each panel show high-magnification views (boxed region) indicating cellular colocalization, with dashed circles marking a representative neuron colocalized with HCAR1 puncta. Scale bar = 50 µm.

We next performed RNAscope to detect HCAR1 expression in p10 hippocampal neurons, a region central to neonatal HI seizures^24,31–33^. RNA scope paired with immunostaining for NeuN (a neuronal marker) shows colocalization of HCAR1 expression and neuronal immunoreactivity (Figure 1B).

### Lactate decreases neuronal excitability and excitation in vitro only in the presence of HCAR1

First, membrane properties were recorded from CA1 principal neurons from p10 C57Bl/6 mice pre and post application of lactate. Lactate hyperpolarized neurons, the mean resting membrane potential was −61.3mV pre-lactate and dropped to −65.1mV post-lactate (n=4 neurons from 2 mice).

Spontaneous excitatory postsynaptic potentials (sEPSC) were recorded from CA1 principal neurons from p10 HCAR1 knockout mice and wildtype littermates before and during application of 3mM of lactate. In wildtype littermates, lactate application shifted the amplitude curve to the left indicating decreased amplitudes after lactate application compared to baseline (p<0.0001, Kolmogorov-Smirnov test, n=6 cells from 3 mice). Conversely, in HCAR1 knockout mice, lactate application did not show any effect on the amplitude curve, indicating no difference in amplitudes after lactate application (p=0.26, Kolmogorov-Smirnov test, n=6 cells from 3 mice). (Figure 2A&B) In addition, interevent interval shifted to the right during lactate application in wildtype mice, indicating decreased sEPSC frequency (p<0.001, Kolmogorov-Smirnov test, n=6 cells from 3 mice) but not in knockout mice (p=0.91, Kolmogorov-Smirnov test, n=6 cells from 3 mice). (Figure 2C)

**Figure 2:**
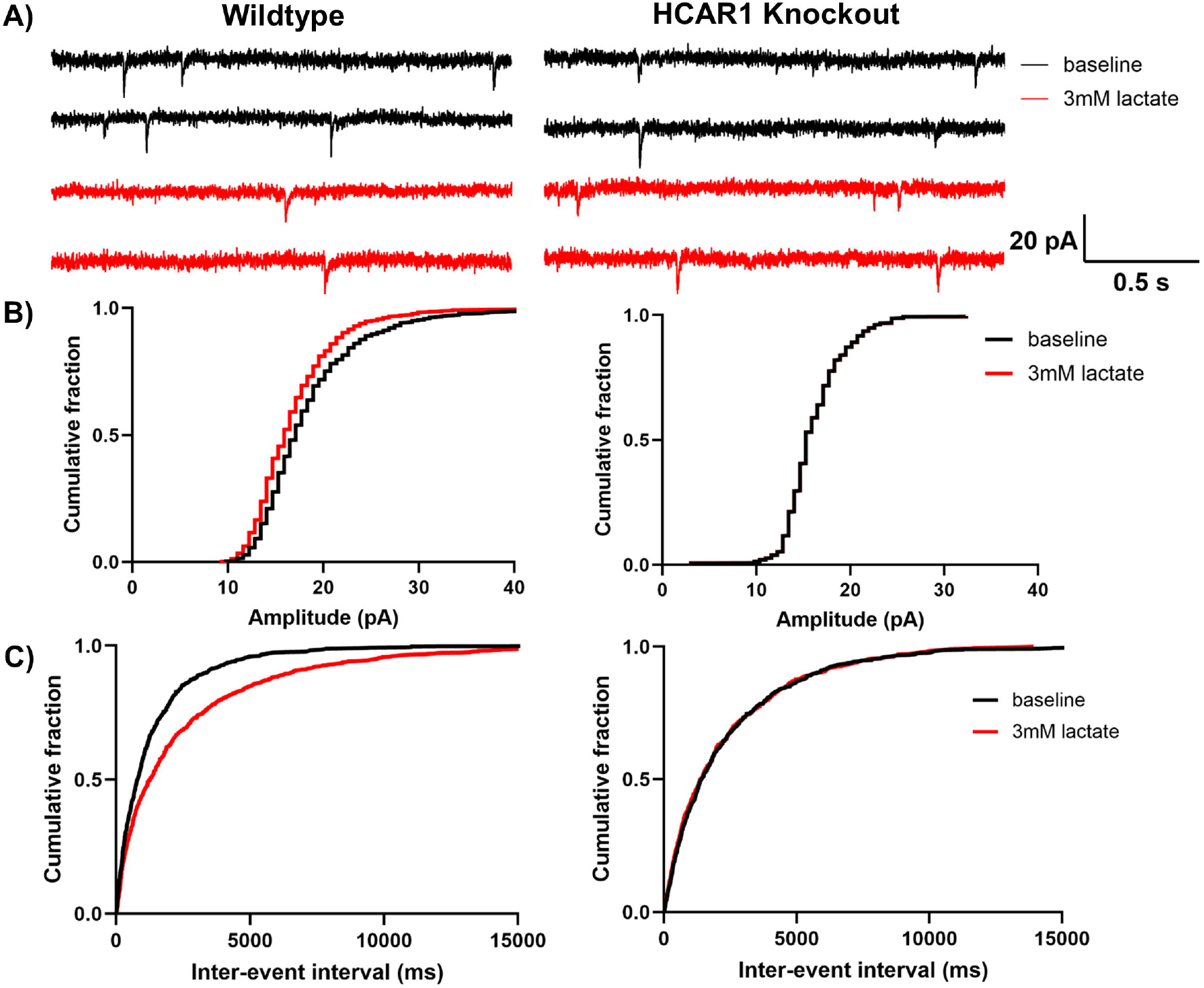
Lactate decreases excitatory transmission in p10 CA1 principal neurons in wildtype but not HCAR1 knockout mice. **A)** Raw traces of sEPSCs in wildtype mice (left) and knockout mice (right) at baseline (black) and after application of 3mM of lactate (red). **B)** Lactate shifts the amplitude curve to the left in WT (left) indicating decreased amplitudes after application of lactate compared to baseline (p<0.0001), lactate did not alter the curve in KO (right) (p=0.26). **C)** Inter-event interval shifted to the right during lactate application in WT (left) indicating decreased sEPSC frequency (p<0.001) but not in KO (right) (p=0.91). n=6 cells from 3 animals per group. Kolmogorov-Smirnov test used in B and C.

### Absence of HCAR1 is associated with increased severity of neonatal HI seizures

HCAR1 knockout mice exhibited an increased number of seizure events over the recording period (87 vs 48 events, n=9/group) and a higher mean seizure duration (61.79±54.43 sec vs. 32.19±26.77 sec, p<0.0001, unpaired t-test) compared to their wildtype littermates (Figure 3A). The HCAR1 knockout mice also exhibited a significantly higher seizure burden over the course of the recording period compared to wildtype mice (6497±5664 sec vs. 2318±3981 sec, p=0.0445, unpaired t-test) (Figure 3B). All seizures on EEG were assigned a behavioral seizure score based on a scale adapted for neonatal rodents^28,34^. HCAR1 knockout mice exhibit more severe behavioral manifestations during seizures with a median BSS of 6 compared to the wildtype group median of 4 (p<0.0001, Mann-Whitney test) (Figure 3C).

**Figure 3:**
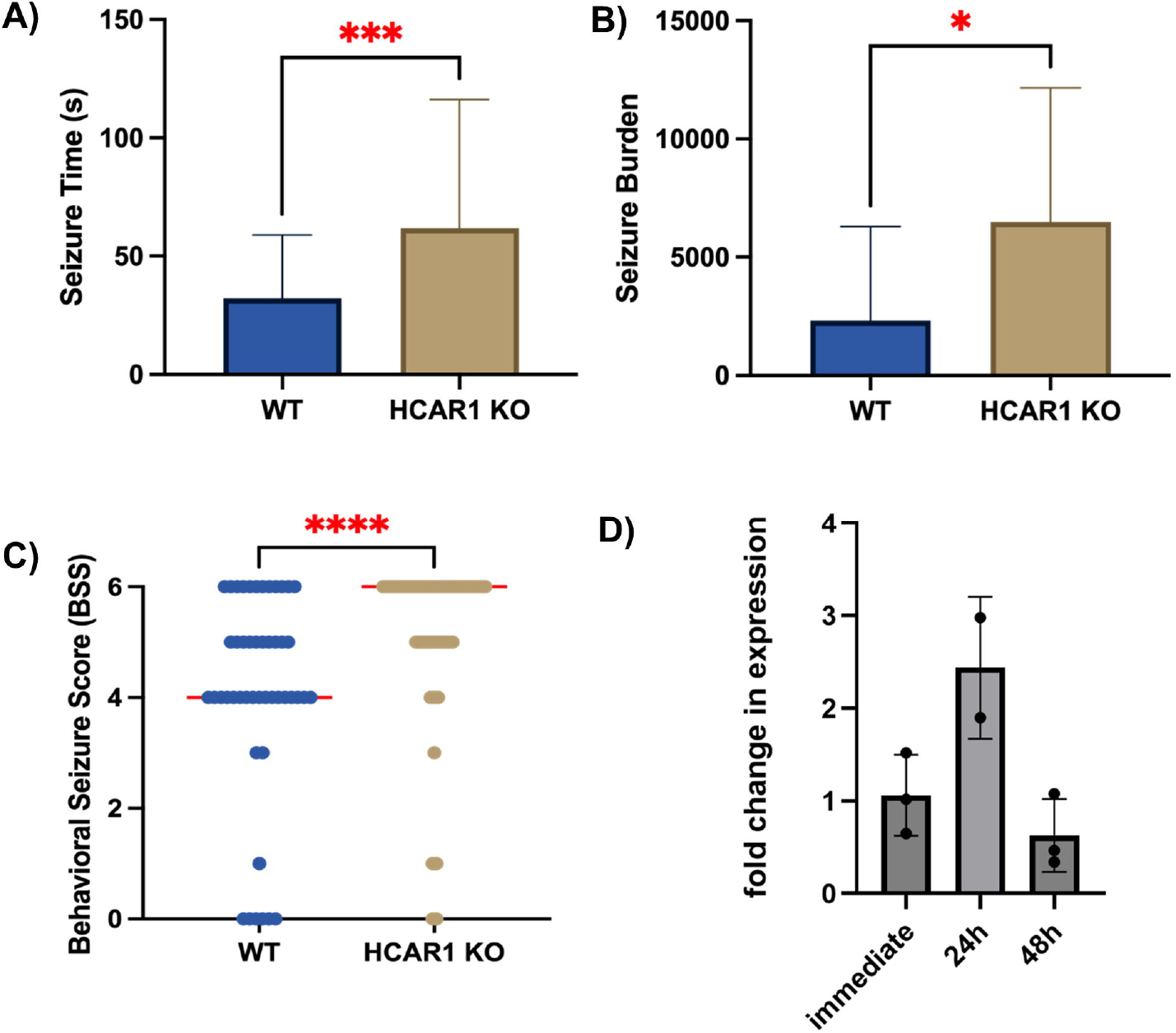
HCAR1 knockout mice have higher seizure burden and severity compared to wildtype during HI and HCAR1 expression is increased post-HI. **A)** Mean seizure duration, 32s in WT vs. 52s in KO (p<0.001). **B)** Seizure burden (#seizures x time per seizure) 2318s in WT vs. 6497s in KO (p=0.04). **C)** Behavioral seizure score median, 4 in WT vs. 6 in KO (p<0.001). (n=9/group); unpaired t-test used in A-C. **D)** In wildtype mice, HCAR1 mRNA expression increased 24h post-HI but drops to below baseline at 48h post-HI. (qPCR, n=3/group; p=0.02, 1-way ANOVA).

### HCAR1 is upregulated in the brain after neonatal HI

Next, we measured and compared HCAR1 mRNA expression in the ipsilateral hemisphere to carotid ligation at three timepoints, immediately, 24 hours after, and 48 hours after HI (Figure 3D). HCAR1 mRNA expression is increased 24 hours post-HI by 2.5x baseline (immediately after HI), then drops to below baseline by 48 hours post-HI (n=3/group; 1-way ANOVA, p=0.027)

## Discussion

In this study we found that HCAR1 is present in the neonatal mouse brain and decreases neuronal excitability *in vitro*. In the context of neonatal hypoxic-ischemic injury, HCAR1 is upregulated after injury and mice lacking HCAR1 exhibit worse acute HI seizures.

Lactate plays a metabolic role in the brain, serving as an alternative energy substrate for neurons. However, lactate also has direct signaling effects in the brain via G-protein coupled receptor, HCAR1^4–9,35^. HCAR1 has been described in adult rodents to play roles in many of the processes central to brain injury and repair, including, neurogenesis, neuronal excitability, angiogenesis, and inflammation^4–8^. Further, HCAR1 is expressed in adult human brain tissue^1–3^.

There is limited data in the neonatal brain regarding HCAR1 expression, function, and its potential role in brain injury. Systemic lactate administration following neonatal HI in a rat model was associated with decreased lesion size^36^ and improved behavioral performance^17,36^. HCAR1’s role in the neuroprotection observed in these studies was unclear, as lactate could be neuroprotective by serving as an alternative metabolic energy source during the energy failure associated with acute HI injury. Both preclinical models and human studies of adult traumatic brain injury (TBI) have demonstrated that administration of exogenous lactate exerts neuroprotection via several mechanisms, including decreasing cerebral edema, functioning as an alternate energy source to glucose, and increasing cerebral blood flow^37–40^. One recent study examined the effect of sodium L-lactate administration in a preterm neonatal population and found that when compared to sodium chloride infusion, sodium L-lactate increased pH and reduced blood lactate levels^41^. However, no studies have examined any neuroprotective effect or other off-target effects of lactate infusion in human neonates and the specific mechanism lactate may play in the neonatal brain remains unclear.

The role and specific mechanisms of HCAR1 signaling in the context of neonatal brain injury remain poorly understood. HCAR1 is expressed in both adult rodent and human brains in neurons and glia^1–3^. HCAR1 activation by a direct agonist decreases neuronal excitability and neuronal network activity^5^. Lactate binding to HCAR1 reduces epileptiform activity in vitro^6^ and lactate produced by excess neuronal activity binds HCAR1 to slow neuronal firing and inhibits excitatory transmission^35^. Further, adult HCAR1 knockout mice exhibit increased seizure severity, duration, and mortality^42^.

Here, we demonstrate that HCAR1 is expressed in neonatal mice at levels similar to the young adult mouse. We also found that HCAR1 is upregulated in the 24 hours following neonatal hypoxia-ischemia. Further, HCAR1 knockout mice exhibit worse acute seizures associated with hypoxia ischemia. Another study found that HCAR1 knockout mice have larger lesion size and lack the normal neurogenesis and gliogenesis found after neonatal hypoxia-ischemia^8^. These findings implicate HCAR1 in both acute injury and tissue repair after neonatal brain injury. The upregulation of HCAR1 in the 24 hours after injury may indicate an ideal therapeutic window for HCAR1 activation to be exploited for neuroprotection. Systemic administration of lactate may have off-target effects and directed HCAR1 signaling with a specific receptor agonist may avoid these effects. Future studies should examine the specific role of HCAR1 direct signaling effects and lactate in neuroprotection after newborn brain injury.

